# Lipoxin A_4_ yields an electrophilic 15-oxo metabolite that mediates FPR2 receptor-independent anti-inflammatory signaling

**DOI:** 10.1101/2024.02.06.579101

**Authors:** Adolf Koudelka, Gregory J. Buchan, Veronika Cechova, James P. O’Brien, Heng Liu, Steven R. Woodcock, Steven J. Mullett, Cheng Zhang, Bruce A. Freeman, Stacy L. Gelhaus

**Author notes:** Correspondence: Bruce A. Freeman; Stacy L. Gelhaus. Greg Buchan, Quest Diagnostics, 6701 Carnegie Avenue, Cleveland, OH 44103, James P. ÒBrien, Polycarbin, 4909 Central Avenue, Richmond, CA 94804.

## Abstract

The enzymatic oxidation of arachidonic acid is proposed to yield trihydroxytetraene species (termed lipoxins) that resolve inflammation via ligand activation of the formyl peptide receptor, FPR2. While cell and murine models activate signaling responses to synthetic lipoxins, primarily 5*S*,6*R*,15*S*-trihydroxy-7*E*,9*E*,11*Z*,13*E*-eicosatetraenoic acid (lipoxin A_4_, LXA_4_), there are expanding concerns about the biological formation, detection and signaling mechanisms ascribed to LXA_4_ and related di- and tri-hydroxy ω-6 and ω-3 fatty acids. Herein, the generation and actions of LXA_4_ and its primary 15-oxo metabolite were assessed in control, LPS-activated and arachidonic acid supplemented RAW 264.7 macrophages. Despite protein expression of all enzymes required for LXA_4_ synthesis, both LXA_4_ and its 15-oxo-LXA_4_ metabolite were undetectable. Moreover, synthetic LXA_4_ and the membrane permeable 15-oxo-LXA_4_ methyl ester that is rapidly de-esterified to 15-oxo-LXA_4_, displayed no ligand activity for the putative LXA_4_ receptor FPR2, as opposed to the FPR2 ligand WKYMVm. Alternatively, 15-oxo-LXA_4_, an electrophilic α,β-unsaturated ketone, alkylates nucleophilic amino acids such as cysteine to modulate redox-sensitive transcriptional regulatory protein and enzyme function. 15-oxo-LXA_4_ activated nuclear factor (erythroid related factor 2)-like 2 (Nrf2)-regulated gene expression of anti-inflammatory and repair genes and inhibited nuclear factor (NF)-κB-regulated pro-inflammatory mediator expression. LXA_4_ did not impact these macrophage anti-inflammatory and repair responses. In summary, these data show an absence of macrophage LXA_4_ formation and receptor-mediated signaling actions. Rather, if LXA_4_ were present in sufficient concentrations, this, and other more abundant mono- and poly-hydroxylated unsaturated fatty acids can be readily oxidized to electrophilic α,β-unsaturated ketone products that modulate the redox-sensitive cysteine proteome via G-protein coupled receptor-independent mechanisms.

## Introduction

Inflammatory responses initiate diverse free radical and enzymatic oxidation reactions of unsaturated fatty acids, yielding a broad array of products that can orchestrate either pathogenic or tissue-protective responses (1–3). A subset of unsaturated di- and tri-hydroxy fatty acid lipid mediators, termed lipoxins, resolvins, maresins and protectins, are products of the enzymatic oxygenation of arachidonic acid (AA, 20:4), eicosapentaenoic acid (EPA, 20:5) and docosahexaenoic acid (DHA, 22:6) (4,5). For the AA-derived lipoxins of focus herein [specifically 5*S*,6*R*,15*S*-trihydroxy-7*E*,9*E*,11*Z*,13*E*-eicosatetraenoic acid; lipoxin A_4_, LXA_4_], trihydroxytetraene formation requires multiple oxygenation reactions catalyzed by cells expressing 5-lipoxgenase (LO), 5-LO-activating protein (FLAP), and 12/15-LO. Two major routes of lipoxin synthesis have been proposed. The first occurs in platelets where leukotriene A_4_ is a substrate for 12-LO and the second involves the action of two lipoxygenases, 5-LO and 12/15-LO, in leukocytes. Based on previous studies it is most likely that 5-LO and FLAP catalyze the first oxygenation of AA, which is then followed by the second oxygenation by 12/15-LO in the same cell or through transcellular synthesis in a neighboring leukocyte (6–9).

The inactivation of LXA_4_ bioactivity is proposed to be the further oxidation of the C15 hydroxyl group by a dehydrogenase such as 15-hydroxy prostaglandin dehydrogenase (15-PGDH), yielding an α, β-unsaturated ketone (10). This class of electrophilic metabolites can come from the diet, intermediary metabolism, fatty acid oxidation and in aggregate are conferred with the ability to regulate inflammatory and metabolic homeostasis through the post translational modification of proteins having reactive nucleophilic amino acids, primarily cysteine (11). The electrophilic moiety of 15-oxo-LXA_4_ can be inactivated by prostaglandin reductase 2 (PTGR2), yielding the non-electrophilic product 13,14-dihydro-15-oxo-LXA_4_ (Fig. 1) (6,12–14).

**Figure 1.**
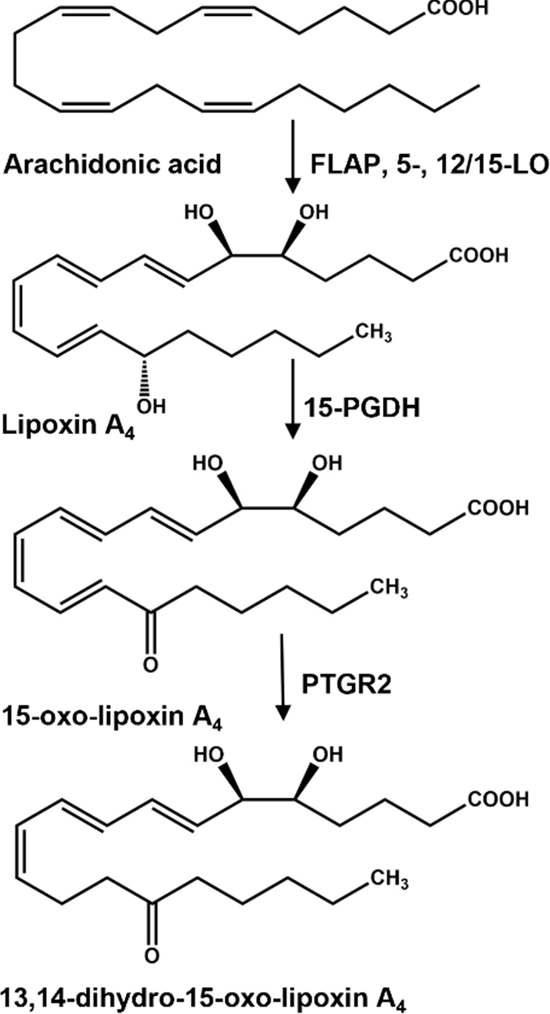
Formation and degradation of lipoxin A_4_. Lipoxin A4 (LXA_4_) is produced from the oxygenation of free arachidonic acid by a combination of 5-lipoxygenase (5-LO), 5-lipoxygenase activating protein (FLAP) and 12/15-lipoxygenase (12/15-LO). LXA_4_ can be further oxidized to an alpha, beta-unsaturated carbonyl containing fatty acid, 15-oxo-LXA_4_, by 15-hydroxyprostaglandin dehydrogenase (15-PGDH).15-oxo-LXA_4_ is further metabolized by prostaglandin reductase 2 (PTGR2) that reduces the C=C bond at C13-C14 rendering the metabolite, 13, 14-dihydro-15-oxo-LXA_4_ non-electrophilic.

Experimental support for the anti-inflammatory and tissue repair-related “specialized pro-resolving mediator” (SPM) actions of trihydroxytetraenes stems exclusively from preclinical studies of synthetic LXA_4_ homologs in biochemical test systems and murine models of inflammatory-related diseases including acute lung injury, asthma, subarachnoid hemorrhage and acute renal failure (8,15–21). Prior to the suggested PTGR2 inactivation of lipoxins, these species are also proposed as specific ligands for the formyl peptide receptor (FPR2), a G-protein coupled receptor (GPCR) that in turn transduces LXA_4_ inhibition of inflammatory signaling and promotion of tissue repair (8,10,22–25). Multiple investigators have raised concerns about ascribing FPR2 as a transducer of lipoxin signaling, due to contradictory observations coming from a) FPR2 knockdown and FPR2 deficient model systems, b) minimal or no changes in cellular Ca^2+^ homeostasis and cyclic AMP levels in test systems treated with synthetic lipoxin homologs, c) limited β-arrestin recruitment to membranes of LXA_4_-activated cells and d) concerns about the stability of LXA_4_ in commercial products and test systems, all of which are comprehensively reviewed (9,26). Because endogenously-generated lipoxins have not been definitively shown to exert specific signaling responses via FPR2 receptor-dependent mechanisms, nor rise to an *in vivo* concentration where they would be expected to do so, alternative explanations may account for the anti-inflammatory and adaptive signaling actions of synthetic lipoxin homologs reported for biochemical, cellular and *in vivo* test systems (6,27).

There are growing numbers of reports contradicting previous data and concepts surrounding the biological generation, endogenous levels, and mechanisms of action of lipoxins and related SPM, initially proposed as endogenous receptor-dependent adaptive signaling mediators. Specifically, multiple labs report undetectable to very low concentrations of SPMs in preclinical and human specimens, even after supplementation with polyunsaturated acids (27–29). In the most rigorous determinations, specimens have been analyzed by mass spectrometry-based methods employing synthetic standards, stable isotope labeled internal standards and the application of internationally-accepted definitions of valid signal to noise responses. These analytical approaches are contrasted by less discriminating approaches to SPM bioanalysis where the identification and quantification of many SPM are based on weak or absent peaks in ion chromatograms and mass spectra that are extracted from sample recordings that are not in concordance with synthetic SPM standard product ion spectra (6,27,30–32).

Multiple reports form the rationale for the present study, including the a) short half-life of most polyunsaturated fatty acid signaling mediators (33–35), b) very low to no detectable tissue, plasma and urine concentrations of trihydroxytetraenes reported *in vivo* (27,36,37), c) appreciation that diverse oxidoreductases such as 15-PGDH and dehydrogenase reductase 9 (SDR9C4) rapidly oxidize hydroxyl derivatives of unsaturated fatty acids to electrophilic α,β-unsaturated ketones (10,38), and d) knowledge that α,β-unsaturated ketone-containing fatty acid derivatives, such as the LXA_4_ metabolite 5,6-dihydroxy-15-oxo-7,9,11,13-eicosatetraenoic acid (15-oxo-LXA_4_), are Michael acceptors and can react with redox-sensing nucleophilic centers of small molecules and proteins to regulate a broad array of electrophile-sensitive transcriptional regulatory and catalytic protein activities, thus impacting downstream signaling (11,39–41). This led to the hypothesis that trihydroxylated unsaturated fatty acids such as LXA_4_ and other related SPMs, if formed in a sufficient concentration *in vivo* or as synthetic pharmacological agents, signal via G-protein coupled receptor-independent electrophilic signaling mechanisms. This hypothesis was based on an appreciation that a) there is a broad spectrum of hydroxylated unsaturated fatty acids are generated by metabolism and inflammatory responses (42,43), and b) the cellular oxidation of hydroxy-fatty acids by enzymes such as 15-PGDH, SDR9C4 and many other dehydrogenases, yields electrophilic α,β-unsaturated ketones that display numerous bioactivities in common with SPM (44,45).

Herein we report that RAW 264.7 macrophages are metabolically competent to mediate LXA_4_ biosynthesis. However, these macrophages did not generate detectable LXA_4_ or its 15-oxo-LXA_4_ metabolite, even when supplemented with AA, activated by LPS and analyzed by liquid chromatography high resolution mass spectrometry (LC-HRMS) methods. Treatment of RAW 264.7 macrophages with 15-oxo-LXA_4_-methyl ester (15-oxo-LXA_4_-Me), a synthetic membrane-permeable precursor for the electrophilic LXA_4_ metabolite 15-oxo-LXA_4_ (46), inhibited NF-ĸB-regulated cytokine expression and promoted nuclear factor (erythroid related factor 2)-like 2 (Nrf2)-dependent adaptive gene expression responses. LXA_4_ did not impact macrophage Nrf2 target gene expression and NF-ĸB-regulated pro-inflammatory mediator expression. Neither LXA_4_ nor its 15-oxo-LXA_4_ metabolite displayed FPR2 ligand activity. It was concluded that multi-target electrophile-mediated signaling may occur through the oxidation of LXA_4_ to 15-oxo-LXA_4_, thus accounting for many of the responses to synthetic LXA_4_ added to *ex vivo* and *in vivo* test systems. The present results motivate further investigation as to whether signaling-competent trihydroxytetraenes occur biologically and if so, whether endogenous concentrations of di- and trihydroxy fatty acids or their electrophilic metabolites are present at levels that would transduce signaling responses via electrophilic metabolite-mediated post-translational modification of redox-sensitive target proteins.

## Results

### RAW 264.7 macrophages express all enzymes needed for the oxidation of AA to LXA_4_

The formation of AA hydroperoxide regioisomers needed for the endogenous synthesis of LXA_4_ are catalyzed by consecutive oxygenation reactions by FLAP and 5-LO, and 12/15-LO (6,12–14). All proteins were detectable in RAW cell lysates over the 24 hr study period (Fig. 2A**-C**). 15-PGDH, which further oxidizes LXA_4_ to 15-oxo-LXA_4_ (44,47), was expressed throughout the 24 hr (Fig. 2D). LPS treatment of RAW macrophages showed the protein expression of FLAP, 5-LO, 12/15-LO and 15-PGDH to be modestly down-regulated (Fig. 2A**-D**). For subsequent studies, an LPS concentration was selected so that there was a maximum of 20% loss of RAW cell viability over 24 hr, an effect that was mitigated by AA supplementation (**Supp.** Fig. 1).

**Figure 2.**
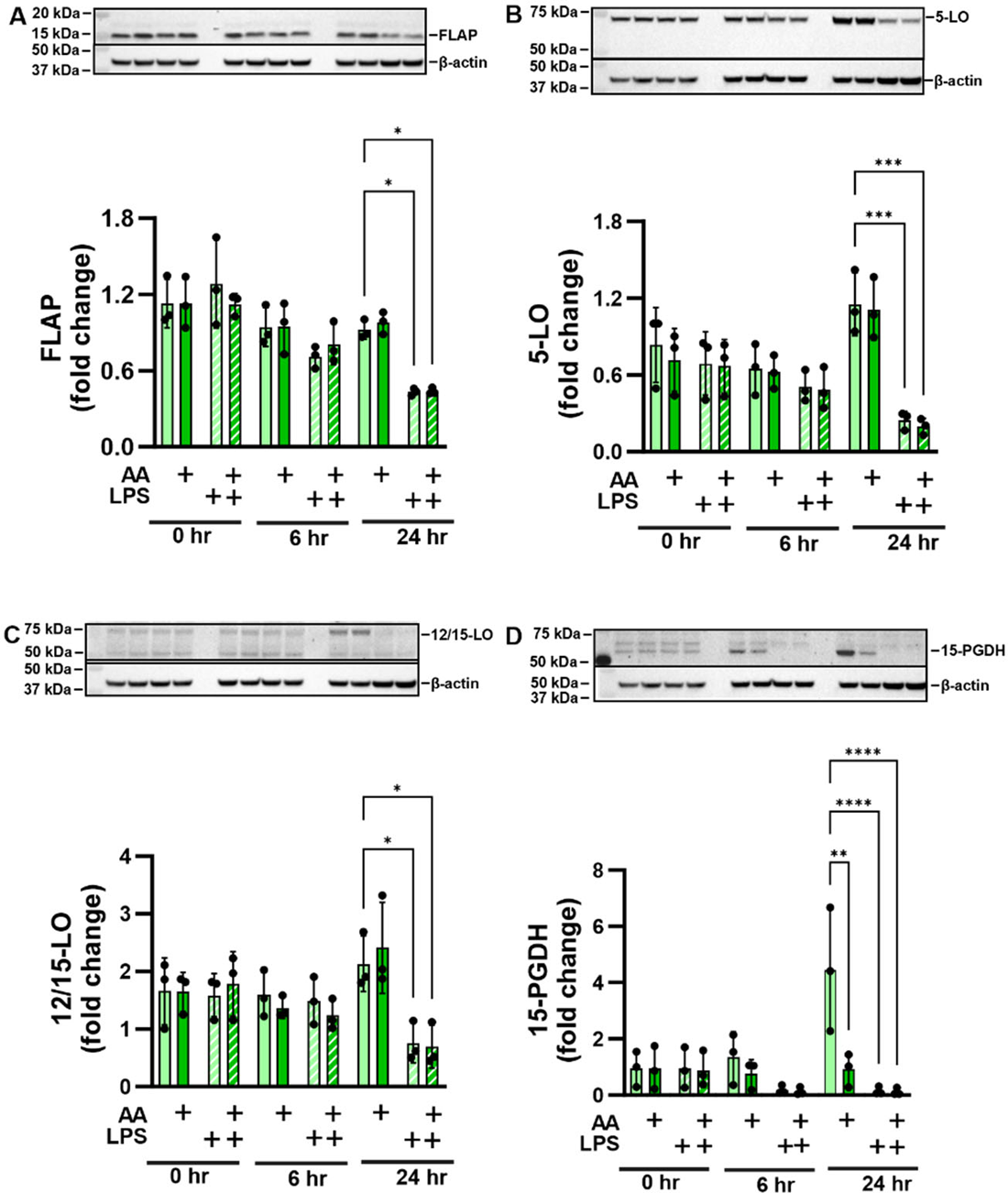
Protein expression of lipoxin synthesis enzymes. Raw 264.7 macrophage express (A) 5-lipoxygenase activating protein (FLAP), (B) 5-lipoxygense (5-LO), (C) 12/15-lipoxygenase (12/15-LO), and (D) 15-hydroxyprostaglandin dehydrogenase (15-PGDH) at basal conditions and upon stimulation with LPS and/or arachidonic acid supplementation over a 24 hr time course.

### LXA_4_ production was not detectable in RAW macrophages under basal, AA and LPS-supplemented conditions

Although all enzymes necessary to produce LXA_4_ were present, LC-HRMS analysis did not show a peak with the matching retention time, accurate mass and MS^2^ fragmentation pattern in the cell lysate (Fig. 3A) or cell media (Fig. 3B), even following supplementation with AA and LPS. In the cell supernatant, a peak with the same accurate mass of LXA_4_ (*m/z* 351.2179, M-H^+^) was observed, but at an earlier retention time of 5.36 min, compared to that of the LXA_4_ standard (**Supp.** Fig. 2A) and the LXA_4_-d_5_ internal standard (*m/z* 355.2494, M-H^+^), both at a retention time of 6.55 min (**Supp.** Fig. 2B). Upon further investigation of the MS^2^, the isobaric species at 5.36 min was likely a prostaglandin metabolite, as evidenced by the characteristic fragment ion of *m/z* 271.2068.

**Figure 3.**
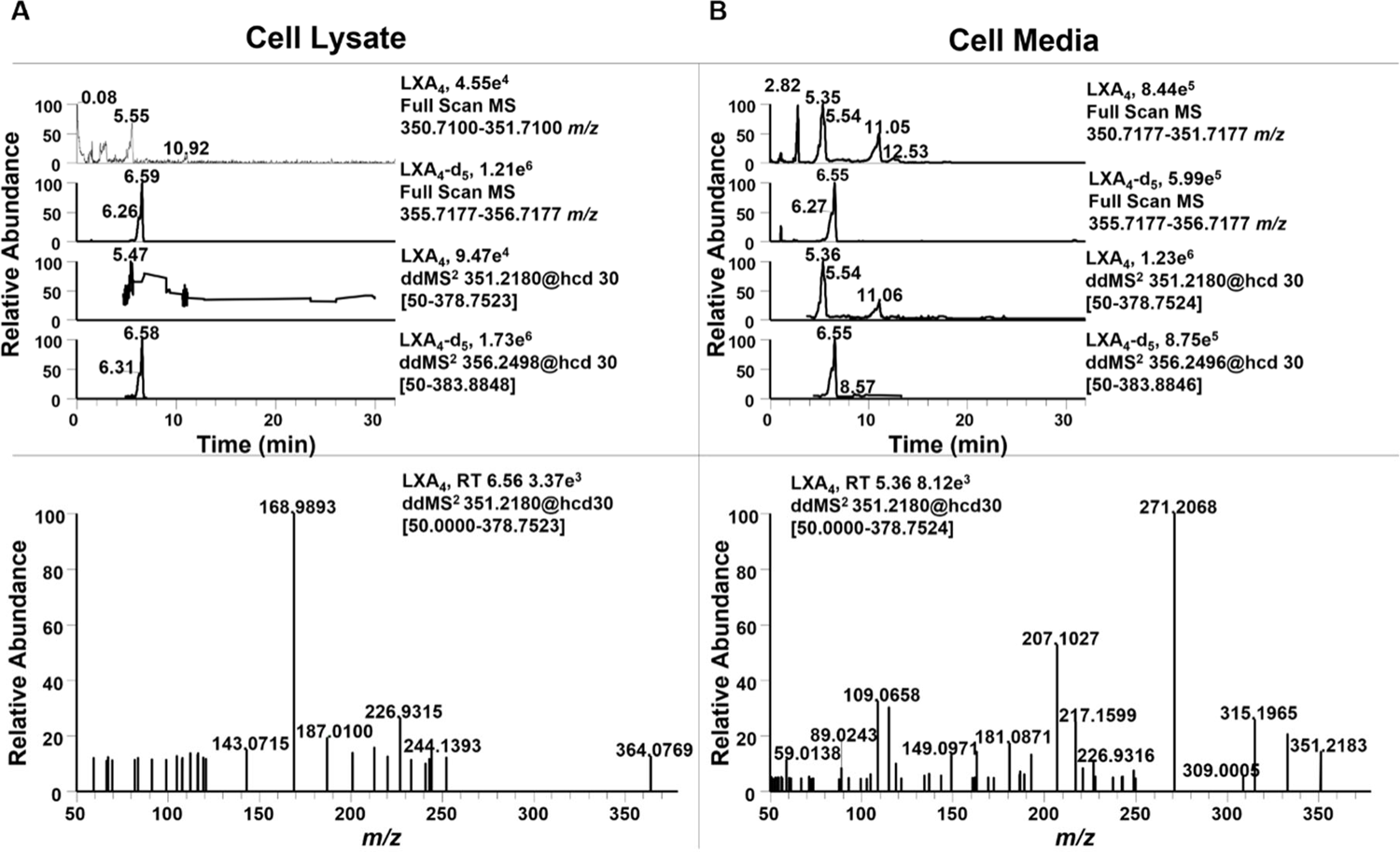
Raw 264.7 macrophage do not produce LXA_4_. Raw 264.7 macrophage were simultaneously supplemented with 25 µM arachidonic acid and stimulated with 10 ng/mL LPS for 24 hr. (A) Cell lysate and (B) Cell supernatant was extracted using a modified Folch extraction containing the internal standard LXA_4_-d_5_ for analysis of oxylipins by LC-HRMS. LXA_4_ was not observed in cell lysate or supernatant. In cell supernatant an oxylipin with the same accurate *m/z* of 351.2183, but different retention time (5.36 min vs 6.55 min) was observed. The MS^2^ for this metabolite contained a diagnostic fragment ion at *m/z* 271.2068 indicating it was a prostaglandin derivative.

### RAW 264.7 macrophages oxidize exogenously-added LXA_4_ to the electrophilic metabolite 15-oxo-LXA_4_

RAW macrophages were treated with 25 µM LXA_4_ over a 12 hr period to investigate the formation of 15-oxo-LXA_4_. LXA_4_ was readily detected in the cell lysate (Fig. 4A), as was 15-oxo-LXA_4_ (**Fig 4B**). Cell lysate levels of LXA_4_ increased early on and remained constant between 3 and 12 hr whereas intracellular levels of 15-oxo-LXA_4_ remained constant. Inversely, LXA_4_ levels remained constant in the media (Fig. 4C) and 15-oxo-LXA_4_ levels increased in the media over the first 6 hr and remained constant between 6 and 12 hr (Fig. 4D). 15-oxo-LXA_4_ eluted at a retention time of 7.28 min with a *m/z* of 349.2028 in both full scan and ddMS^2^ mode (Fig. 4E). To confirm the identity of 15-oxo-LXA_4_, cells were treated with 15-oxo-LXA_4_-Me that is rapidly de-esterified by methyl transferases (46). A peak having a retention time 7.32 min was detected for the de-esterified 15-oxo-LXA_4_-Me product, with a parent *m/z* of 349.2020 for the M-H^+^ ion (Fig. 4F). The product ion spectra for 15-oxo-LXA_4_ and the de-esterified 15-oxo-LXA_4_-Me standard showed matching diagnostic ions at *m/z* 331.1919, 287.2019, 233.1548, 189.1285, 165.1284, 139.1128, 113.0971, and 69.0345 (Fig. 4G and 4H).

**Figure 4.**
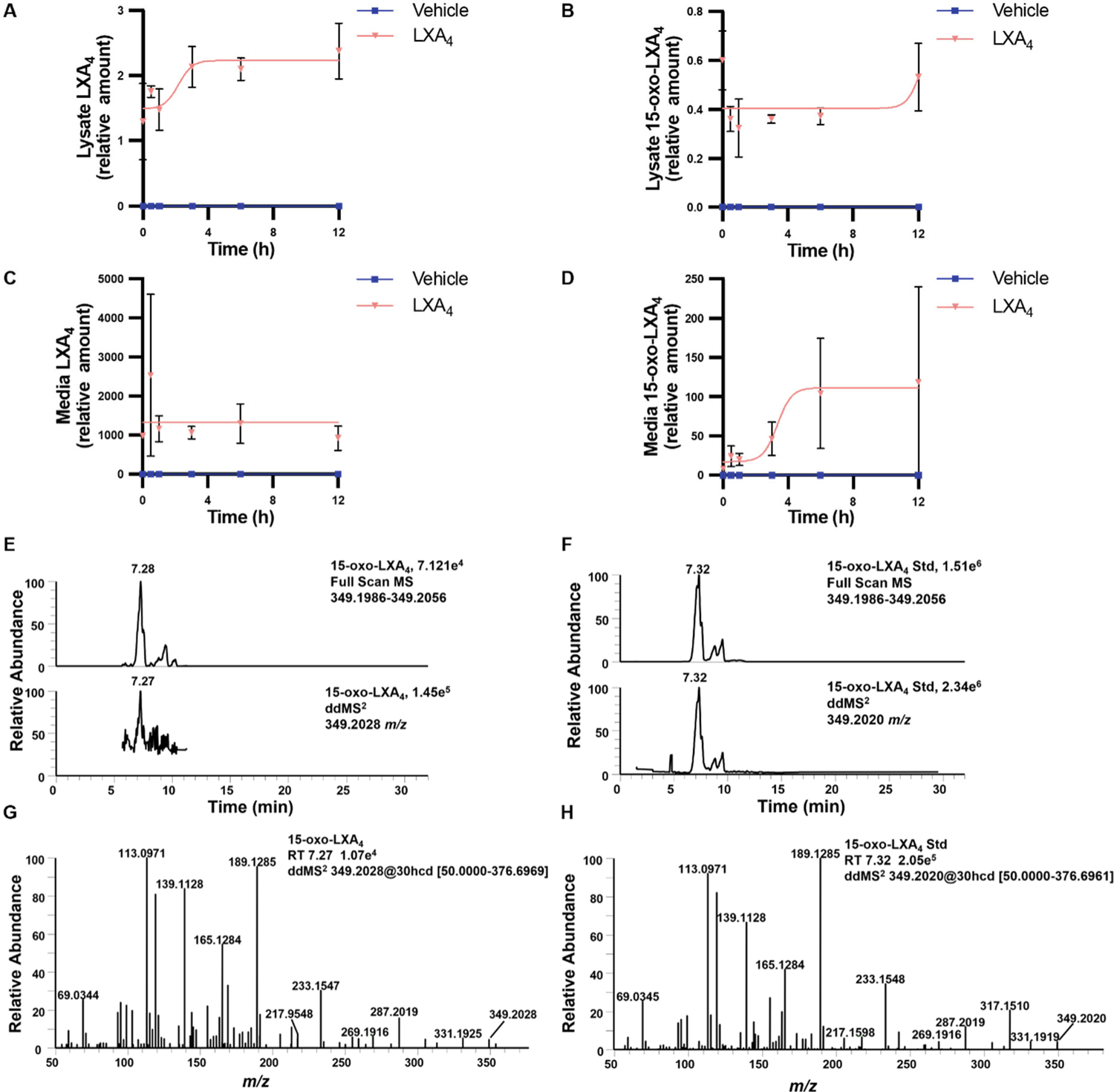
Raw 264.7 macrophage metabolize exogenous LXA_4_ to 15-oxo-LXA_4_. Raw 264.7 macrophage were treated with 25 µM LXA_4_ over a 12 hr time course and (A) LXA_4_ and (B) 15-oxo-LXA_4_ were measured in cell lysate and (C) LXA_4_ and (D) 15-oxo-LXA_4_ were measured in cell media. (E) The endogenously formed 15-oxo-LXA_4_ was retained at 7.28 min and was detected in both full scan and ddMS^2^ with a *m/z* of 349.2028. (F) 15-oxo-LXA_4_-Me was added to Raw 264.7 cells for 1 hr to create an ionizable standard for comparison of retention time and product ion MS^2^ and displayed matching *m/z* at 349.2020 at a retention time of 7.32 min. The product ion spectra of the (G) the endogenous 15-oxo-LXA_4_ analyte and (H) the 15-oxo-LXA_4_ standard have corresponding diagnostic fragment ions at *m/z* 331.1925, 233.1547, 189.1285, 165.1284, 139.1128, and 113.0971.

### Signaling actions of LXA_4_ and 15-oxo-LXA_4_-Me are FPR2-independent

The FPR2 receptor is ligand-activated by the formylated peptide fMet-Leu-Phe and related fMet peptides, annexin-1 and amyloid-α and-β. FPR2 is the proposed receptor for LXA_4_ and other oxygenated unsaturated fatty acids (24,38). Herein, the FPR2 receptor was expressed by RAW macrophages over the course of the study period, with cell protein expression levels not affected by AA supplementation and LPS treatment (Fig. 5A). A sensitive *in vitro* GTPγS binding assay was performed to investigate the ligand-induced activation of FPR2 (48,49). The assay measures the binding of ^35^S-GTPγS to Gi, the cognate signaling partner of FPR2, as an indicator of FPR2 activation. The FPR2 peptide agonist, WKYMVm (50,51) was used as a positive control, with FPR2 activation inducing significant ^35^S-GTPγS binding to Gi. In contrast, LXA_4_ and its metabolite 15-oxo-LXA_4_ did not display any FPR2 receptor agonism at any concentration tested (Fig. 5B).

**Figure 5.**
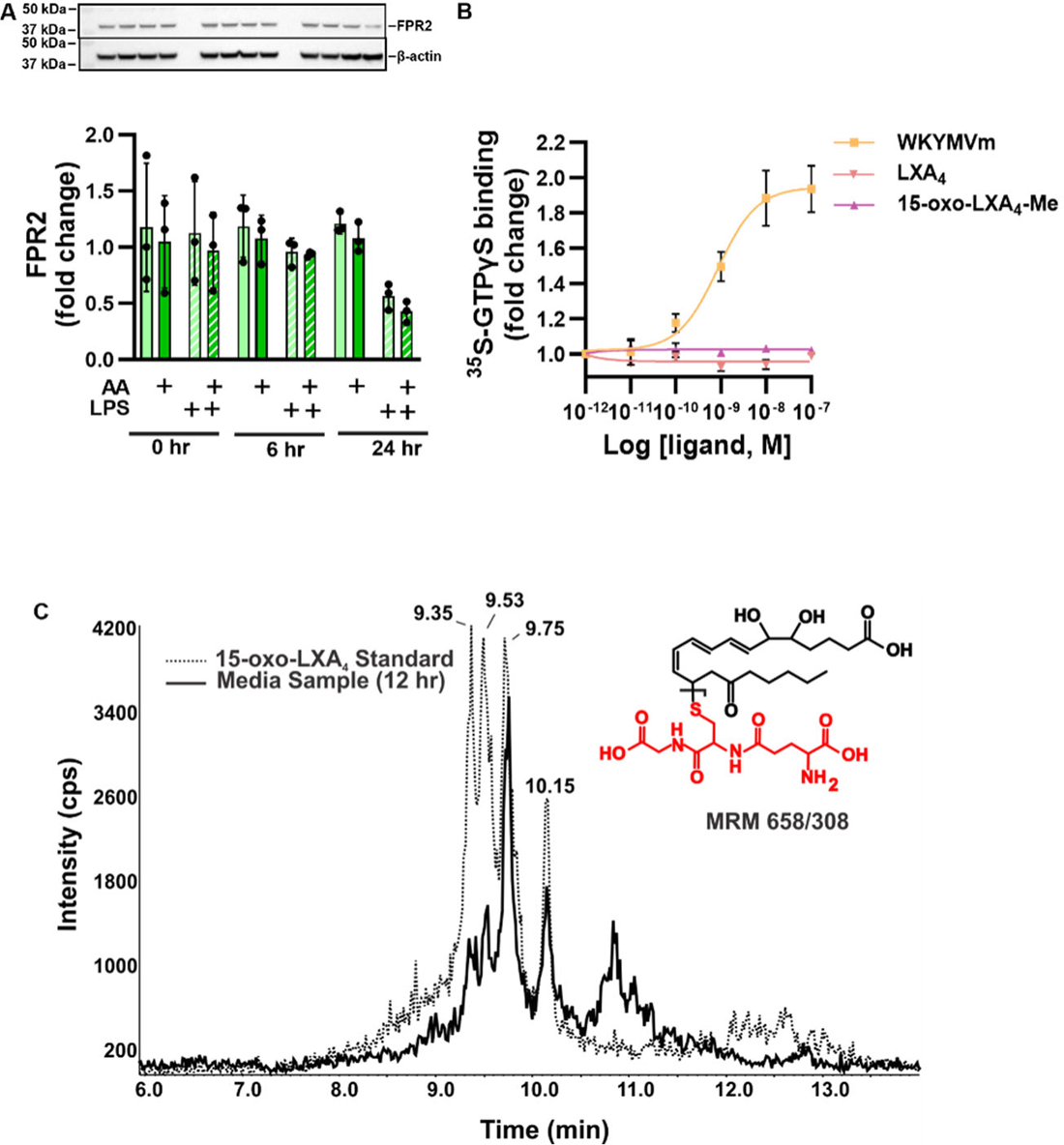
LXA_4_ and 15-oxo-LXA_4_-Me do not activate FPR2. (A) FPR2 expression was measured by western blot in Raw 264.7 macrophage. (B) Increasing concentrations of LXA_4_, 15-oxo-LXA_4_ and the FPR2 ligand WKYMVm were incubated with S-GTPγ and binding was determined. (C) Chromatographic overlay of glutathionylated 15-oxo-LXA_4_ standard and 15-oxo-LXA_4_ glutathione adduct in Raw cell media at 12 hr after treatment with 15-oxo-LXA_4_-Me.

To define whether 15-oxo-LXA_4_ is electrophilic and might signal via FPR2-independent mechanisms, 100 µM glutathione and 10 µM 15-oxo-LXA_4_-Me were incubated in 50 mM potassium phosphate buffer (pH = 8) for 1 hr at 37 °C. Glutathionylated 15-oxo-LXA_4_ was monitored by LC-MS/MS in positive ion mode using selected reaction monitoring at the *m/z* transition 658 →308. The GSH adduct was found in cell supernatant of RAW cells treated with 25 µM 15-oxo-LXA_4_-Me. A representative chromatographic overlay is shown in Fig. 5C.

### 15-oxo-LXA4-Me inhibits pro-inflammatory signaling

RAW macrophages were activated with LPS and the activation of NF-ĸB target gene expression was reflected by the increased expression of interleukin (IL)-1β, IL-6, monocyte chemoattractant protein 1 (MCP-1), and inducible nitric oxide synthase (iNOS, Nos2), compared to vehicle controls (Fig. 6A**-D**). There was upregulation of IL-6 and MCP-1 protein expression measured by ELISA (Fig. 6E **and F**) and iNOS by western blot analysis (Fig. 6G). The effects of 15-oxo-LXA_4_-Me on LPS-induced gene expression of pro-inflammatory mediators was variable and primarily occurred at the 12 and 18 hr timepoints. LPS-induced IL-1β mRNA expression was downregulated by 15-oxo-LXA_4_-Me at all measured times, becoming significant at 12-18 hr (Fig. 6A). IL-6 mRNA expression and cytokine production was inhibited at most time points, with statistically significant decreases at 18-24 hr after LPS plus 15-oxo-LXA_4_-Me treatment (Fig. 6B and 6E). *Ccl2* (MCP-1) mRNA expression and cytokine production were significantly inhibited by 15-oxo-LXA_4_-Me 12 hr post LPS administration (Fig. 6C **and F**). 15-oxo-LXA_4_-Me also significantly inhibited LPS-stimulated *Nos2* mRNA expression between 12 and 24 hr (Fig. 6D). Compared with vehicle-treated cells, the maximum loss of viability due to LPS treatment was ∼20%, an effect that was mitigated by 15-oxo-LXA_4_-Me over the 24 hr period. 15-oxo-LXA_4_-Me supplementation alone did not significantly impact cell viability (**Supp.** Fig. 1). The signaling actions of LXA_4_ were compared with 15-oxo-LXA_4_-Me in LPS treated macrophage at the 12 hr time point. For the cytokines IL-1β and MCP-1 (Fig. 6H **and I**) and iNOS protein expression (Fig. 6J) LXA_4_ did not significantly impact the expression of pro-inflammatory signaling mediators when compared to cells treated with 15-oxo-LXA_4_-Me.

**Figure 6.**
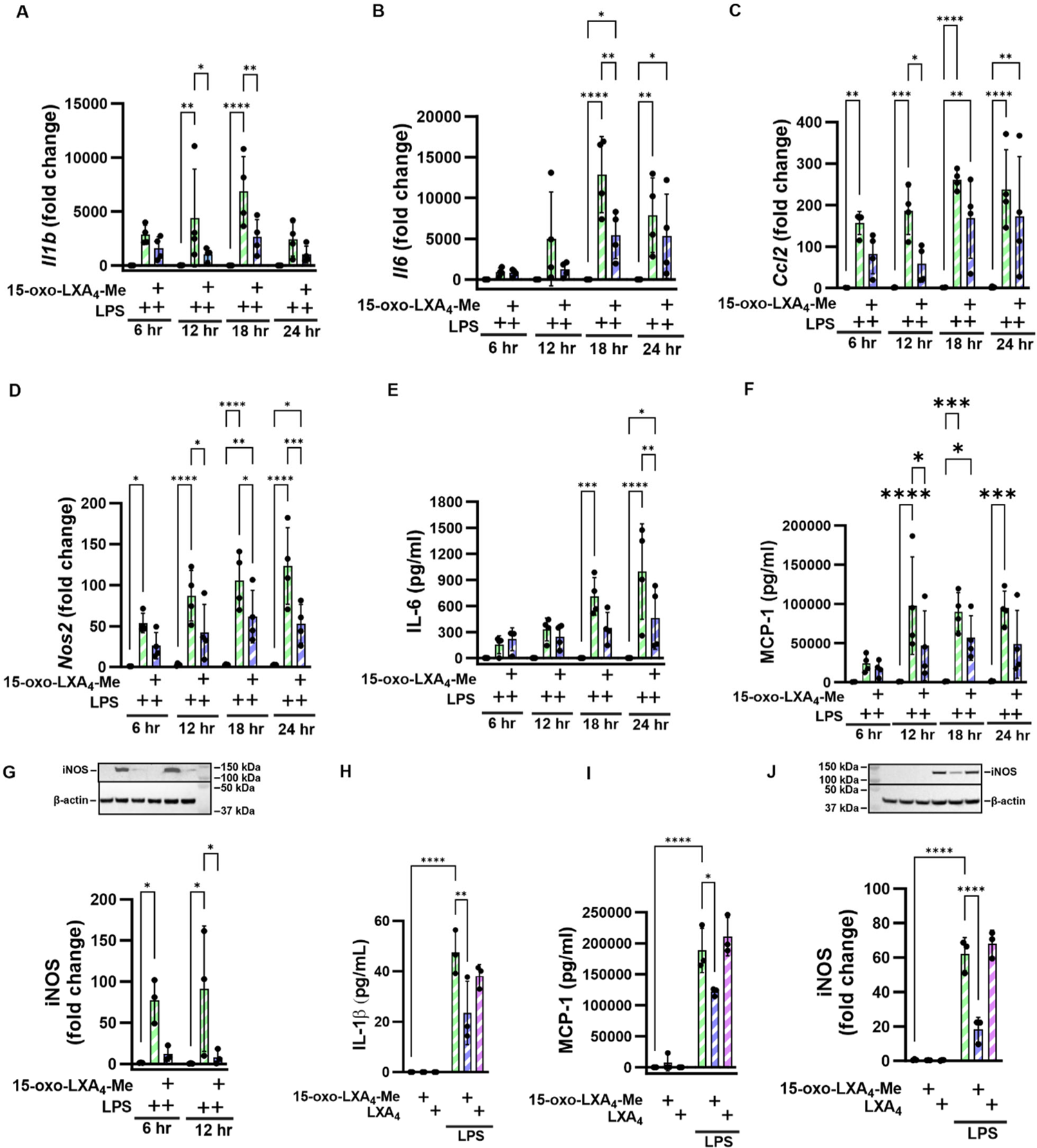
15-oxo-LXA_4_-Me inhibits pro-inflammatory pathways. Treatment of Raw 264.7 macrophage with 15-oxo-LXA_4_-Me inhibits LPS-mediated upregulation of pro-inflammatory signaling across a 24 hr period. (A) *Il1b* (B) *Il6* (C) *Ccl2* (D) *Nos2* gene expression, (E) IL-6 protein (F) MCP-1 protein and (G) iNOS protein expression. At 12 hr 15-oxo-LXA_4_-Me and LXA_4_ inhibition of LPS-mediated upregulation of inflammatory pathways was compared for (H) IL-1β protein (I) MCP-1 protein and (J) iNOS protein. * p < 0.05, ** p < 0.01, *** p < 0.001, **** p < 0.0001.

### 15-oxo-LXA_4_-Me induces anti-inflammatory signaling

The activation of Nrf2-regulated gene and protein expression by 15-oxo-LXA_4_-Me was reflected by the increased expression of its targets including glutamate-cysteine ligase modifier subunit (GCLM), heme oxygenase 1 (HO-1), and NAD(P)H quinone oxidoreductase 1 (NQO1). 15-oxo-LXA_4_-Me increased both gene and protein expression of the Nrf2 regulated targets (Fig. 7A**-F**). *Gclm* and *Ho1* mRNA expression peaked at 6 hr (Fig. 7A **and C**), while the increase in *Nqo1* expression at 6 hr was sustained at 12 hr (Fig. 7E). Western blotting of the Nrf2 target proteins further validated the activation of this electrophile-sensitive transcriptional regulatory mechanism by 15-oxo-LXA_4_-Me at 6 and 12 hr for GCLM and HO-1 (Fig. 7B **and D**), and at 12 hr for NQO1 (Fig. 7F). In contrast to 15-oxo-LXA_4_-Me, LXA_4_ had no significant impact on Nrf2-regulated adaptive signaling responses, both with and without RAW 264.7 cell activation by LPS (Fig. 7G**-I**).

**Figure 7.**
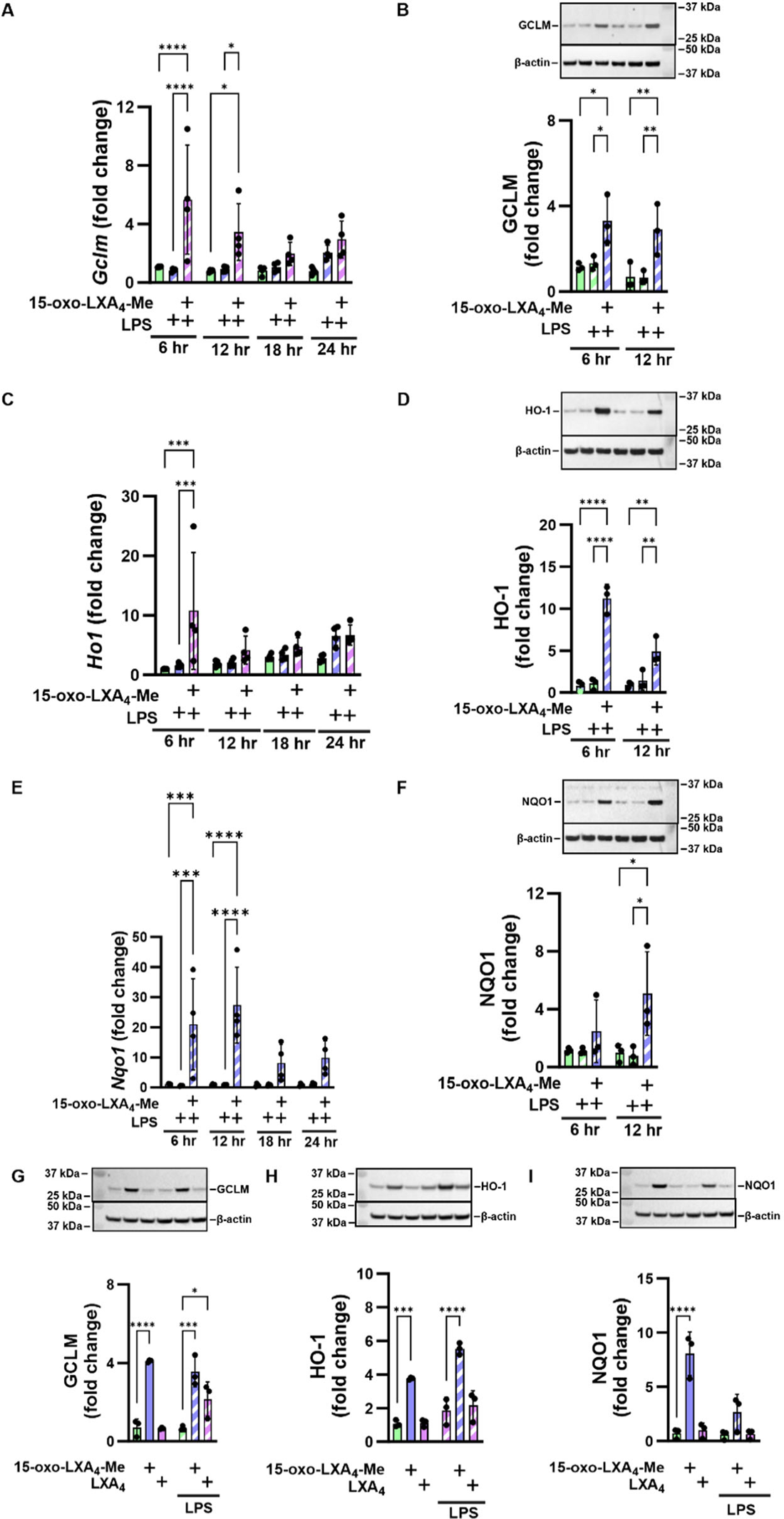
15-oxo-LXA_4_-Me activates the antioxidant response. Treatment of Raw 264.7 macrophage with 15-oxo-LXA_4_-Me results in activation of the antioxidant response after LPS treatment across a 24 hr period. (A) *Gclm* (B) GCLM protein (C) *Ho1* (D) HO-1 protein (E) *Nqo1* (F) NQO1 protein. At 12 hr 15-oxo-LXA_4_-Me and LXA_4_ activation of the antioxidant response after LPS stimulation was compared for (G) GCLM protein (H) HO-1 protein and (I) NQO1 protein. * p < 0.05, ** p < 0.01, *** p < 0.001, **** p < 0.0001.

## Discussion

The formation of di- and tri-hydroxytetraene fatty acids in leukocytes has been studied by various groups in biochemical systems, cell culture and *in vivo* systems since the 1980s (22,23,52). Although many cells express the enzymes required to synthesize lipoxins, the formation of these trihydroxy species are at very low or undetectable endogenous concentrations and that these concentrations are not definitively linked with the resolution of inflammation (6,53–57). Notably, the initial reports of cellular lipoxin A_4_ generation required neutrophil priming with the 15-lipoxygenase product 15-hydroperoxy-5,8,11,13-eicosatetraenoic acid (15-HPETE) to accomplish lipoxin biosynthesis (22,58).

Herein, western blot analysis affirmed the expression of 5-, 12/15-LO, and FLAP protein that would be required for lipoxin biosynthesis in RAW 264.7 macrophages (Fig. 2); however, LXA_4_ generation was undetectable, even with cell activation with LPS and supplementation of AA (Fig. 3). Only upon supplementation of macrophages with synthetic LXA_4_ was the 15-PGDH metabolite, 15-oxo-LXA_4_, detected both intra- and extracellularly (Fig. 4). Issues regarding endogenous LXA_4_ concentrations have gained attention recently, as some investigators are proposing to measure SPM levels to predict clinical and therapeutic outcomes (59–61). Multiple labs, including the present report focusing on 15-oxo-LXA_4_ as an electrophilic signaling mediator, are unable to replicate the biological detection of sufficient concentrations of trihydroxytetraene and other SPM to concentrations that might induce downstream signaling responses *in vitro* and *in vivo* (9,27,36,37). This dilemma is magnified by a) the failure of the primary labs reporting biological SPM generation by LC-MS/MS to apply standard limit-of-detection or limit-of-quantitation criteria to SPM analyses and b) the use of misleading representative LC-MS chromatograms of SPM generation that are not derived from directly-related primary data (37). This has resulted in the triumph of hope over reality in many SPM-related studies, where a flawed characterization of background noise was the criterion for the presence and concentrations of SPM in biological samples. This conundrum motivated the reassessment of past evidence for the presence of SPM in biological matrices and the inferred roles of the SPM LXA_4_ in the resolution of inflammation.

Herein, FPR2 receptor-independent reactions of 15-oxo-LXA_4_-Me are evidenced by the engagement of critical constituents of the redox-sensitive cysteine proteome (NF-ĸB and Keap1/Nrf2) and a lack of FPR2 receptor agonism by both LXA_4_ and 15-oxo-LXA_4_ (**Figs. 5-7**). Current dogma holds that trihydroxytetraene oxidation to the 15-oxo metabolite by 15-PGDH inactivates FPR2 ligand activity and downstream signaling; however, this is not the case. Both LXA_4_ and 15-oxo-LXA_4_-Me displayed a complete lack of ligand-induced activation of FPR2, as measured by GTPγS binding analysis. This contrasted with the robust activation of FPR2-GTPγS binding induced by WKYMVm, a synthetic peptide mimic of N-formylated bacterial peptides, (Fig. 5B). This does not exclude the possibility that both lipids may act on FPR2 to induce the activation of other FPR2 signaling partners such as β-arrestins, again if produced in sufficient concentration (4,10,38,62,63). The pharmacokinetics (PK) of 15-oxo-LXA_4_, if present in sufficient concentrations, will distinctly differ from, and might offer advantages over its precursor, LXA_4_. Following oxidation of LXA_4_ to 15-oxo-LXA_4_, there will be an equilibrium between free and nucleophile-adducted 15-oxo-LXA_4_, as for other small molecule electrophiles. Over time, if LXA_4_ and 15-oxo-LXA_4_ generation is sufficient and continues, 15-oxo-LXA_4_-target protein adducts can accumulate to induce downstream signaling responses from even low rates of LXA_4_ generation (64). Previous PK interpretations of LXA_4_ signaling relies on the weak GPCR ligand activity of LXA_4_, where Michaelis-Menten kinetics will dictate cellular responses. This characteristic could explain how the administration of supra-physiological concentrations of synthetic LXA_4_, other SPMs or their prodrugs might contribute to the propagation of adaptive anti-inflammatory signaling *in vivo*.

Diverse electrophilic unsaturated fatty acids are generated by both enzymatic catalysis and as products of the free radical and oxidizing species that are produced during digestion, basal metabolism and inflammatory responses (43,65). These fatty acid species most typically have electron-withdrawing keto, nitro or halogen substituents on or in conjugation with an alkene (42,66,67). At biological concentrations fatty acid electrophiles form Michael addition adducts, primarily with cysteine and to a lesser extent histidine containing peptides and proteins (65,68–70). This cysteinome includes a) cysteines involved in maintaining the 3-dimensional structure of proteins, b) metabolic and transport enzymes, c) signaling proteins such as phosphatases and kinases, d) proteins that regulate cell growth and differentiation and e) transcriptional regulatory factors that modulate the expression of ∼1% of the human genome (71–79). RNAseq, in concert with functional enrichment analysis affirms that small molecule electrophiles, such as those derived from fatty acids, can mediate the transcriptional regulation of ∼100-250 genes related to metabolism, transport, cell cycle, genome stability, mitochondrion and endoplasmic reticulum organization, proteostasis and inflammation (76,77,79–81). These evolutionarily-conserved transcriptional regulatory mechanisms, that rely on the reactions of a limited number of hyperreactive and functionally-significant nucleophilic amino acids, provides organisms with a capability to respond to increased concentrations of oxidizing and electrophilic species generated during metabolism and inflammation (41,82). In addition to transcriptional regulatory proteins, other proteins that participate in mediating inflammatory and tissue repair responses are also targets of unsaturated fatty acid-derived electrophiles. This includes the nuclear lipid peroxisome proliferator-activated receptor γ (PPARγ), NADPH oxidase-2, xanthine oxidoreductase, cyclooxygenase-2, soluble epoxide hydrolase, diverse protein kinases and phosphatases, matrix metalloproteinase-7 and −9, stimulator of interferon gamma (STING) and calcineurin (83–90).

The generation and metabolism of hydroxyeicosatetraenoic acids (HETEs) is a well-established example of oxidation-induced functional switching of a hydroxy-fatty acid derivative to an α, β-unsaturated ketone product. Following peroxidase reduction of primary unsaturated fatty acid dioxygenation products to hydroxyl moieties (e.g., 5-, 11-, 12- or 15-HETE), 15-PGDH and other cellular dehydrogenases having broad substrate specificities generate α, β-unsaturated ketone products from HETEs. For example, the oxidation of 11-HETE and 15-HETE by 15-PGDH yields 11-oxo-ETE and 15-oxo-ETE, respectively (91–93). Both electrophilic metabolites promote adaptive anti-inflammatory responses by inhibiting NF-ĸB signaling and pro-inflammatory cytokine expression in both murine and human macrophages (91,93). Electrophilic metabolites of hydroxyl moieties in 20 carbon trihydroxypentadienes and 22 carbon trihydroxyhexadienes have been reported both *in vitro* and *in vivo.* For example, 8-oxo-resolvin D1 and 17-oxo-resolvin D1 are formed from resolvin D1 and 18-oxo-resolvin E1 from resolvin E1 (94,95). These metabolites, originally viewed to have lost ligand activity for their cognate GPR32, GPR18 and/or ChemR23 receptors, have yet to be characterized in the context of likely downstream signaling actions expected for the corresponding electrophilic α, β-unsaturated ketone-containing metabolites, an exercise motivated by data presented herein (19,94).

In summary, the electrophilic metabolite of LXA_4_, 15-oxo-LXA_4_, activated Nrf2-regulated antioxidant and tissue repair responses and inhibited LPS-induced, NF-ĸB-regulated pro-inflammatory cytokine expression in RAW 264.7 macrophages. Many of the SPM properties attributed to LXA_4_ and other mono-, di- and trihydroxy fatty acids overlap with those reported for electrophilic metabolites. These results indicate that the LXA_4_ metabolite 15-oxo-LXA_4_ and other more abundant electrophilic omega-3 fatty acid mono-hydroxy derivatives can exert electrophile-responsive, FPR2 receptor-independent effects after further oxidation. For this property to be of biological significance however, LXA_4_ must be generated at sufficient concentrations. The extensive literature describing the multi-target signaling actions of lipid electrophiles indicates that many, if not all of the products of non-enzymatic and enzymatically-catalyzed fatty acid oxidation reactions will induce antioxidant and anti-inflammatory responses upon alkylation of critical nucleophilic amino acids of enzymes and transcriptional regulatory proteins (43,71,72). The present data motivates that the concentrations, metabolism, gene expression responses and the temporal differentiation of GPCR-dependent and GPCR-independent responses to unsaturated fatty acid hydroxyl and oxo fatty acid metabolite levels be examined in more detail.

## Experimental procedures

### Cell cultivation and treatment

RAW 264.7 (ATCC: TIB-71) murine macrophages were grown in DMEM (Corning: 10-013-CV) with 10% FBS (Gibco: 26140-079) and 1% Penicillin/Streptomycin (P/S, Gibco: 15140-122), and used for all studies. RAW cells were plated at 2×10^6^ cells/well in 6 well plates and cultured for 12 hr in DMEM+10% FBS, next media were replaced with DMEM+1% FBS+1% P/S. Cells were treated with LPS (10 ng/mL, Sigma: L4391, Lot: 067M4036V), AA (25 µM, Cayman: 90010), LXA_4_ (25 µM, Cayman: 90410), or 15-oxo-LXA_4_-Me (25 µM) which was synthesized as previously reported (46). Cytotoxicity was determined by MTT [3-(4,5-dimethylthiazol-2-yl)-2,5-diphenyltetrazolium bromide (ThermoFisher: M6494) reduction analysis according to manufacturer’s instructions. Cell lysate and media were collected for endpoint analyses including ELISA, western blot, PCR and liquid chromatography high resolution mass spectrometry (LC-HRMS).

### Western blot

For western blot analysis, cells were scraped into ice cold RIPA buffer (Cell Signaling: 9806) with protease (Roche: 04693132001) and phosphatase (Roche: 4906845001) inhibitors. Samples were further lysed via sonication (10 sec on 5 sec off, repeat 3x, ThermoFisher: FB120, 4°C) and protein was clarified by centrifugation at 21,000 x g for 5 min at 4°C. Protein concentrations were measured using the BCA protein assay (ThermoFisher: 23225) according to manufacturer’s instructions, samples were diluted to 2 mg/mL, and mixed with NuPage sample buffer (ThermoFisher: NP0007) and reducing agent (ThermoFisher: NP0009). Samples were heated at 100°C for 10 min, 20 μL of each sample was loaded onto a polyacrylamide gel (ThermoFisher 4-12%, WG1401BOX and WG1402BOX), 5 μL of dual color standard (Bio-Rad: 1610374) was loaded for molecular weight estimation, and electrophoresis was performed for 1-2 hr at 130 V. Protein was transferred to nitrocellulose membrane (Bio-Rad: 1620115) at 100 V for 1 hr at 4°C. Membranes were washed with TBS-T buffer and blocked with either 5% milk diluted in TBS-T or 1% casein in TBS (ThermoFisher: 37532) for 1 hr at room temperature. Membranes were washed in TBS-T and primary antibodies COX-2 (Cell Signaling: 12282T), FLAP (Invitrogen: PA5-78368), FPR2 (Invitrogen: 720293), GCLM (Invitrogen: PA5-26111), HO-1 (Enzo Life Sciences: ADI-SPA-895-F), iNOS (Cell Signaling: 13120), NQO1 (Abcam: ab80588), 5-LO (Cell signaling: 3289S), 12/15-LO (Invitrogen: MA5-25891), 15-PGDH (Santa Cruz: SC-271418) were added overnight at 4°C. The following day, primary antibodies were washed with TBS-T, and HRP-linked anti-rabbit (Cell Signaling: 7074) or anti-mouse (Cell Signaling: 7076) secondary antibodies were added. Images were taken using ECL substrates (Bio-Rad: 1705061) and a Bio-Rad ChemiDoc imager. Optical density was evaluated in Image Lab software (Bio-Rad). Optical density of β-actin (Sigma: A4700) was used as endogenous control. Representative images of all presented western blots presented for each figure.

### RT-PCR

For PCR analysis, cells were scraped into TRIzol (Invitrogen: 15596026). RNA was isolated, concentration measured, and cDNA prepared as described previously (96). FAM-dyed primers (all purchased from Taqman) were used: *Gclm* (Mm01324400_m1), *Ho1* (Mm00516005_m1), *Il1b* (Mm00434228_m1), *Il6* (Mm99999064_m1), *Ccl2* (Mm00441242_m1), *Nos2* (Mm00440502_m1), *Nqo1* (Mm01253561_m1), and VIC-dyed *Actin* (Mm00607939_s1) was used as endogenous control.

### ELISA

Media were centrifuged at 500 x g for 5 min at 4°C and diluted so that absorbance results were in the range of the standard curve. Specific protocols for each cytokine came from the Invitrogen kits: IL1-β (88-7013-88), IL-6 (88-7064-88), and MCP-1 (Ccl2, 88-7391-88).

### LC-MS Sample Preparation

Media were collected and cells washed 3x with PBS, cells were scraped into PBS and immediately frozen at −80°C. LXA_4_-d_5_ (10 µL of 1 µg/mL stock, manufacturer: cat. #10007737) was added to 1 mL of cell lysate or media. To each sample, 4 mL of chloroform:methanol (2:1) was added, vortexed and centrifuged at 2500 rpm for 10 min at 4°C. The organic layer was dried under nitrogen and solvated in 100 µL methanol on the day of analysis.

*LC-HRMS* Samples were analyzed on a Thermo Fisher Exploris 240 hybrid mass spectrometer coupled to an Vanquish Horizon UHPLC. Samples were applied to a Luna C8 column (2 x100 mm, Phenomenex, cat. 00D-4248-B0) and eluted with a linear gradient using H_2_O with 0.1% acetic acid as solvent A and acetonitrile with 0.1% acetic acid as solvent B. Samples were loaded at 35% B and the gradient increased to 60% B over 30 min, held at 100% B for 2 min, and equilibrated at 35% B for 3 min at a flow rate of 0.3 mL/min. LXA_4_ and LXA_4_ metabolites were measured using negative electrospray ionization under the following MS conditions: source, 2600 V; sheath gas 50, auxiliary gas 10, sweep gas 1, ion transfer temperature 325 °C, and vaporizer temperature 350 °C. Relative levels of LXA_4_ and 15-oxo-LXA_4_ were quantified by normalizing to LXA_4_-d_5_.

*15-oxo-LXA_4_ GSH adduct standard and LC-MS Analysis* A glutathionylated 15-oxo-LXA_4_ standard was made by incubation of 10 µM 15-oxo-LXA_4_-Me with 100 µM GSH in 50 mM potassium phosphate buffer (pH = 8) for 1 hr at 37°C (91,97). GSH conjugates were extracted from 1 mL of cell supernatant using Oasis HLB 1 cc solid phase extraction columns (Waters) and applied to a Luna C18 column (2 x 100 mm, Phenomenex) at a flow rate of 0.25 mL/min and eluted with a linear consisting of solvent A (H2O + 0.1% acetic acid) and Solvent B (ACN + 0.1% acetic acid). The gradient started at 20% B at 5 min and increasing to 98% B at 25 min. The gradient was held at 100% B for 2 min and equilibrated at 20% B for 35 min. GSH adducts were analyzed on a 6500+ QTRAP coupled to an Exion LC (Sciex) using multiple reaction monitoring (658→308) and positive ionization with the following MS conditions: CUR 40, CAD med, IS 4500, GS1 70, GS2 65, Temp 550°C, DP 80, EP 7, CE 17, CXP 7.

### GTPγS binding assay

The GTPγS binding assay was conducted using cell membranes overexpressing FPR2 and purified G_i_ heterotrimer, following our previously reported method (49). Briefly, Sf9 insect cells (ExpressionSystems) were infected with baculovirus expressing FPR2. The cells were collected by centrifugation at 8000 x g for 10 min after 48 hr. For ^35^S-GTPγS binding analysis, ∼200 µg/mL of human FPR2 cell membrane was incubated with 200 nM purified G_i_ protein for 20 min on ice in buffer containing 20 mM HEPES, pH 7.5, 150 mM NaCl, 5 mM MgCl_2_, 3 μg/mL BSA, 0.1 μM TCEP, and 5 μM GDP. Next, 25 μL FPR2-G_i_ mix was transferred to 225 μL reaction buffer containing 20 mM HEPES, pH 7.5, 150 mM NaCl, 5mM MgCl_2_, 3 μg/mL BSA, 0.1 μM TCEP, 1 μM GDP, 35 pM ^35^S-GTPγS (Perkin Elmer) and ligands (LXA_4_ and 15-oxo-LXA_4_-Me at 2 μM, WKYMVm (Tocris) at 5 μM). After additional 15 min incubation at 25°C, the reaction was terminated by adding 6 mL of cold wash buffer containing 20 mM HEPES, pH 7.5, 150 mM NaCl and 5mM MgCl_2_, and filtering through glass fiber prefilters (Millipore Sigma,). After washing 4 times with 6 mL cold wash buffer, the filters were incubated with 5 mL of CytoScint liquid scintillation cocktail (MP Biomedicals) and counted on a Beckman LS6500 scintillation counter.

*Statistical analysis* was performed using GraphPad Prism 9 (GraphPad Software). Bar graph data represent mean values of all results +/- SD, with each individual result represented by black dot. All data presented in the bar graphs were tested via 2-way analysis of variance. Symbols in XY graphs represent mean values of all results +/- SD. Data were tested via nonlinear regression (curve fit), represented by connecting curve. Data represents 3-4 independent experiments. Non-statistically significant results are denoted as ns, meaning p > 0.05, all statistically significant results are denoted by asterisks, with * p ≤ 0.05, ** p ≤ 0.01, *** p ≤ 0.001, **** p ≤ 0.0001.

## Author Contributions

AK, GJB, SLG, SRW, JPO, VC, SJM, HL, and CZ conducted experimental work. AK, GJB, CZ, BAF, and SLG designed and analyzed experiments and then prepared the manuscript draft approved by all contributors.

## Acknowledgements

This work was supported by NIH T32GM008424 and F31HL142171 (GJB), R35GM128641 (CZ), R01 HL132550 and P01 HL103455 (BAF), S10OD032141 (SLG), and R33HL157069 (BAF, SLG). We would like to acknowledge Larry Marnett and Valerie O’Donnell for their helpful discussions.

## Additional Information

BAF acknowledges an interest in Creegh Pharmaceuticals.

