## Supplemental Figures 1 and 2 for "Lipoxin A_4_ yields an electrophilic 15-oxo metabolite that mediates FPR2 receptor-independent anti-inflammatory signaling"

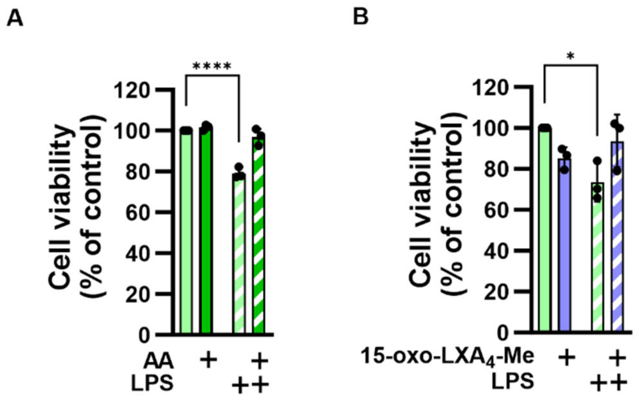


**Supplemental Figure 1. Raw 264.7 macrophage viability after LPS, AA and 15-oxo-LXA$_4$-Me treatment**. Raw 264.7 macrophage were treated for 24 hr with (A) 10 ng/mL LPS, 10 μM AA, or in combination and (B) 10 ng/mL LPS, 25 μM 15-oxo-LXA$_4$-Me, or in combination. Cell viability was measured using the MTT assay. * $p < 0.05$ and **** $p < 0.0001$.

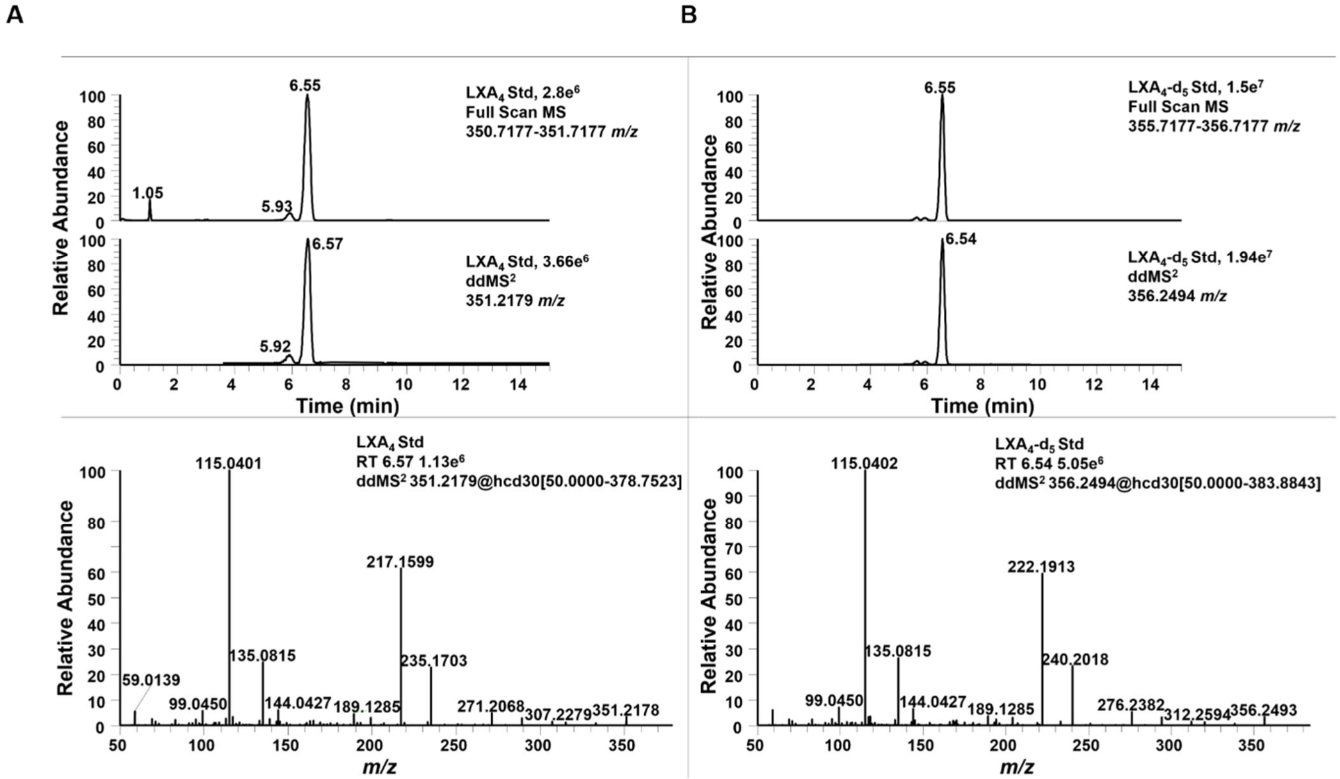


**Supplemental Figure 2. LC-HRMS of $LXA_4$ and $LXA_4$-$d_5$ commercial standards**. (A) The top panel shows the chromatogram for $LXA_4$ with a retention time of 6.55 min in both full scan and $ddMS^2$ modes. The bottom panel shows the product ion spectra for *m/z* 351.2179 at 6.55 min. Diagnostic ions for $LXA_4$ include *m/z* 307.2279, 271.2068, 235.1703, 217.1599, 135.0815, 115.0401. (B) The top panel shows the chromatogram for $LXA_4$-$d_5$ with a retention of 6.55 min in both full scan and $ddMS^2$. The bottom panel shows the product ion spectra for *m/z* 356.2494 at 6.55 min. Diagnostic ions for $LXA_4$-$d_5$ include *m/z* 312.2594, 276.2382, 240.2018, 222.1913, 189.1285, 135.0815, and 115.0402.
